# TE invasion fuels molecular adaptation in laboratory populations of *Drosophila melanogaster*

**DOI:** 10.1101/2022.06.06.494973

**Authors:** Luyang Wang, Shuo Zhang, Savana Hadjipanteli, Lorissa Saiz, Lisa Nguyen, Efren Silva, Erin S. Kelleher

## Abstract

Transposable elements are mobile genetic parasites that frequently invade new host genomes through horizontal transfer. Invading TEs often exhibit a burst of transposition, followed by reduced transposition rates as repression evolves in the host. We recreated the horizontal transfer of *P*-element DNA transposons into a *D. melanogaster* host, and followed the expansion of TE copies and evolution of host repression in replicate laboratory populations reared at different temperatures. We observed that while populations maintained at high temperatures rapidly go extinct after TE invasion, those maintained at lower temperatures persist, allowing for TE spread and the evolution of host repression. We also surprisingly discovered that invaded populations experienced recurrent insertion of *P*-elements into a specific long non-coding RNA, *lncRNA:CR43651*, and that these insertion alleles are segregating at unusually high frequency in experimental populations, indicative of positive selection. We propose that, in addition to driving the evolution of repression, transpositional bursts of invading TEs can drive molecular adaptation.

## Introduction

Transposable elements (TEs) are mobile genetic parasites that spread through host populations by producing new chromosomal insertions that are transmitted from parent to offspring. Despite this system of vertical transmission, comparative genomics increasingly reveals that TEs are frequently transferred horizontally between non-mating species, introducing new TEs into the host genome (Gilbert et al. 2010; Thomas et al. 2010; Reiss et al. 2019). After these invasions, TE copy numbers often increase rapidly owing to the absence of host regulation of the novel parasite (Pace et al. 2008; Kofler et al. 2015).

Invading TE families have significant impacts on the fitness of the host. Unregulated transposition results in a high frequency of novel and predominantly deleterious mutations (Slatko and Hiraizumi 1973; Robertson 1978). Additionally, in sexually reproducing organisms, DNA damage resulting from transposition can disrupt the developmental process of gametogenesis, reducing fecundity and blocking the progression of embryogenesis (Schaefer et al. 1979; Orsi et al. 2010; Renaut and Bernatchez 2011). However, the initial impact of TE invasions on host populations, as well as subsequent host adaptation, are challenging to study because they are brief selective events that have often occurred in the distant past. Additionally, host repression may evolve rapidly after invasion, thereby masking the impacts of unregulated transposition (Kofler et al. 2018; Zhang et al. 2020).

In the last 100 years, multiple *Drosophila* species’ genomes have been invaded by *P*-element DNA transposons, providing snapshots of how TE invasion impacts the host, and how the host responds. *P*-elements were introduced into *D. melanogaster* from its distant relative *D. willistoni* (MRCA ~40 MYA) in the 1950s (Kidwell 1983; Anxolabéhère et al. 1988). They subsequently spread to *D. melanogaster* populations worldwide, into *D. simulans* populations worldwide (Kofler et al. 2015), and a recent report suggests an even more recent invasion into *D. yakuba* (Serrato-Capuchina et al. 2018) In *D. melanogaster* and *D. simulans, P*-element invasion has been connected with a sterility syndrome known as hybrid dysgenesis, which reduces or prevents the production of viable gametes (Kidwell et al. 1977; Hill et al. 2016; Serrato-Capuchina et al. 2018). Furthermore, in *D. melanogaster* and *D. simulans* host regulation of *P*-elements evolved rapidly, within <30 years after invasion (Kidwell 1983; Hill et al. 2016).

*P*-element repression in natural populations of *D. simulans* and *D. melanogaster* occurs through the production of Piwi-interacting RNAs (piRNAs) that are maternally transmitted from parent to offspring (Brennecke et al. 2008; Hill et al. 2016; Zhang et al. 2020). The production of repressive piRNAs from antisense *P*-element transcripts is initiated when *P*-element insertions occur in piRNA clusters: the genomic loci that produce precursor piRNAs (Ronsseray et al. 1996; Marin et al. 2000; Stuart et al. 2002; Brennecke et al. 2008; Kofler et al. 2018; Zhang et al. 2020). Interestingly, while *P*-element insertion alleles confer a profound increase in host fertility in the presence of genomic *P*-elements, they do not show clear signatures of recent positive selection in natural populations of *D. melanogaste*r (Zhang et al. 2020). This apparent paradox is partially explained by the presence of recombination, which rapidly separates repressor alleles from chromosomes they have protected from transposon load, thereby limiting their selective advantage (Charlesworth and Langley 1986). In addition, the neutral evolution of repression is facilitated by the frequent production of repressor alleles through new insertions into piRNA clusters, which avoids a requirement for strong natural selection (Kelleher et al. 2018; Kofler 2019; Zhang et al. 2020).

To recapture the impacts of *P*-element invasion on host populations and further dissect the evolutionary response of the host, we recreated the invasion in laboratory populations over a three year period. We combined time-course phenotyping of hybrid dysgenesis and piRNA production to reveal the fertility costs of invasion and detect the emergence of repression by the host. We further employed deep sequencing of pooled population samples to reveal the propagation of *P*-elements throughout the genome, as well as the occurrence of insertions into piRNA producing sites. Consistent with previous work from ourselves and others, we observed that piRNA-mediated silencing evolves rapidly after invasion via the preferential transposition of *P*-elements into piRNA clusters located in telomere associated sequences (TAS) (Kofler et al. 2018; Zhang et al. 2020). Unexpectedly however, we also discovered biased transposition of *P*-elements into a non-piRNA producing long non-coding RNA, *lncRNA:CR43651*. These insertions occurred recurrently in independent laboratory populations, and exhibited unusually high allele frequencies. Our observations suggest that while piRNA cluster insertions may not be strongly beneficial, invading TEs may promote adaptation in their host by producing adaptive insertions in other loci.

## Results

We recreated the *P*-element invasion by introducing full-length active *P*-elements into a naive genotype (KSA2/A4) via germline transformation. 10 replicate populations were created from independent transformation of the KSA2 background, and maintained simultaneously at low (22°C) and high temperatures (27°C). To detect the expansion of genomic *P*-elements and their repression by piRNAs, we phenotyped these populations for induction and repression of hybrid dysgenesis every 5 generations for 51 generations, or until the population went extinct (Figure 1). The capacity of experimental males to paternally induce F1 hybrid dysgenesis in crosses with naive females (*P*-element free, M strain) is correlated with the paternal abundance of genomic *P*-elements (Bingham et al. 1982; Srivastav and Kelleher 2017; Serrato-Capuchina et al. 2020). By contrast, the ability of experimental females to maternally repress F1 hybrid dysgenesis in crosses with inducer (P strain) males that harbor abundant genomic *P*-elements is indicative of evolved piRNA-mediated repression (Brennecke et al. 2008).

**Figure 1.**
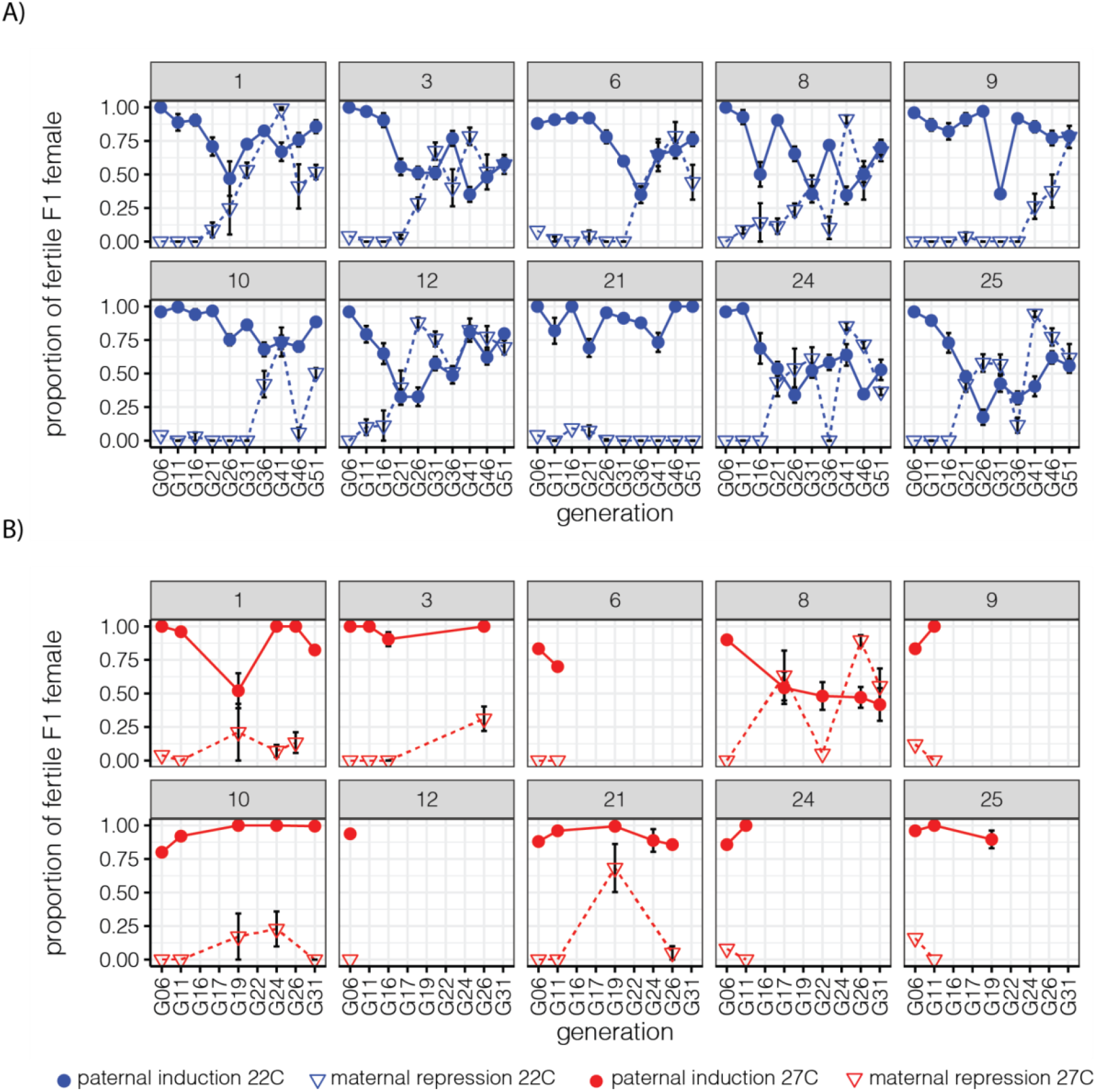
Phenotypic evolution in laboratory populations. Induction and repression of *P*-element hybrid dysgenesis was estimated every 5 generations by quantifying F1 ovarian atrophy in test crosses. Induction is indicated by filled circles and repression is indicated by triangles. Error bars correspond to standard error in the estimated proportion, based on replicate crosses involving individual experimental males or females. A) 22 °C. B) 27°C.

### Rapid evolution of induction and repression in colonized populations maintained at 22°C

With one exception (Population 21), females from laboratory populations exhibited a systematic increase in maternal repression of hybrid dysgenesis, suggesting the evolution of piRNA mediated silencing (Figure 1A, Supplementary Table 1, Supplementary Table 2). Similarly, males from 7 out 10 populations exhibited increased induction of hybrid dysgenesis over the course of the experiment, while two populations (12 and 21) exhibited a decrease in induction, and one population (1) exhibited no systematic change (Figure 1A, Supplementary Table 1, Supplementary Table 2).

To contrast the genomic invasion of *P*-elements among populations we deep sequenced 20 Generation 51 females from each of the 22°C populations (411-650X coverage, Supplementary Table 3), yielding an average coverage of ~10-32X per haploid genome. With the exception of population 21, populations reared at 22°C contained 9-15 *P*-element copies per haploid genome based on sequencing coverage (Figure 2A), and 21-29 copies per haploid genome based on the summed frequency of individual insertion alleles (Figure 2B). The higher copy numbers resulting from allele frequency-based estimates likely results from increased read mapping when aligning to individual insertions. In population 21, individuals were estimated to contain far fewer *P*-element copies (Figure 2A-B, .39 copies based on coverage, .76 copies based on allele frequency), which is consistent with the absence of induction and repression over the course of the experiment.

**Figure 2.**
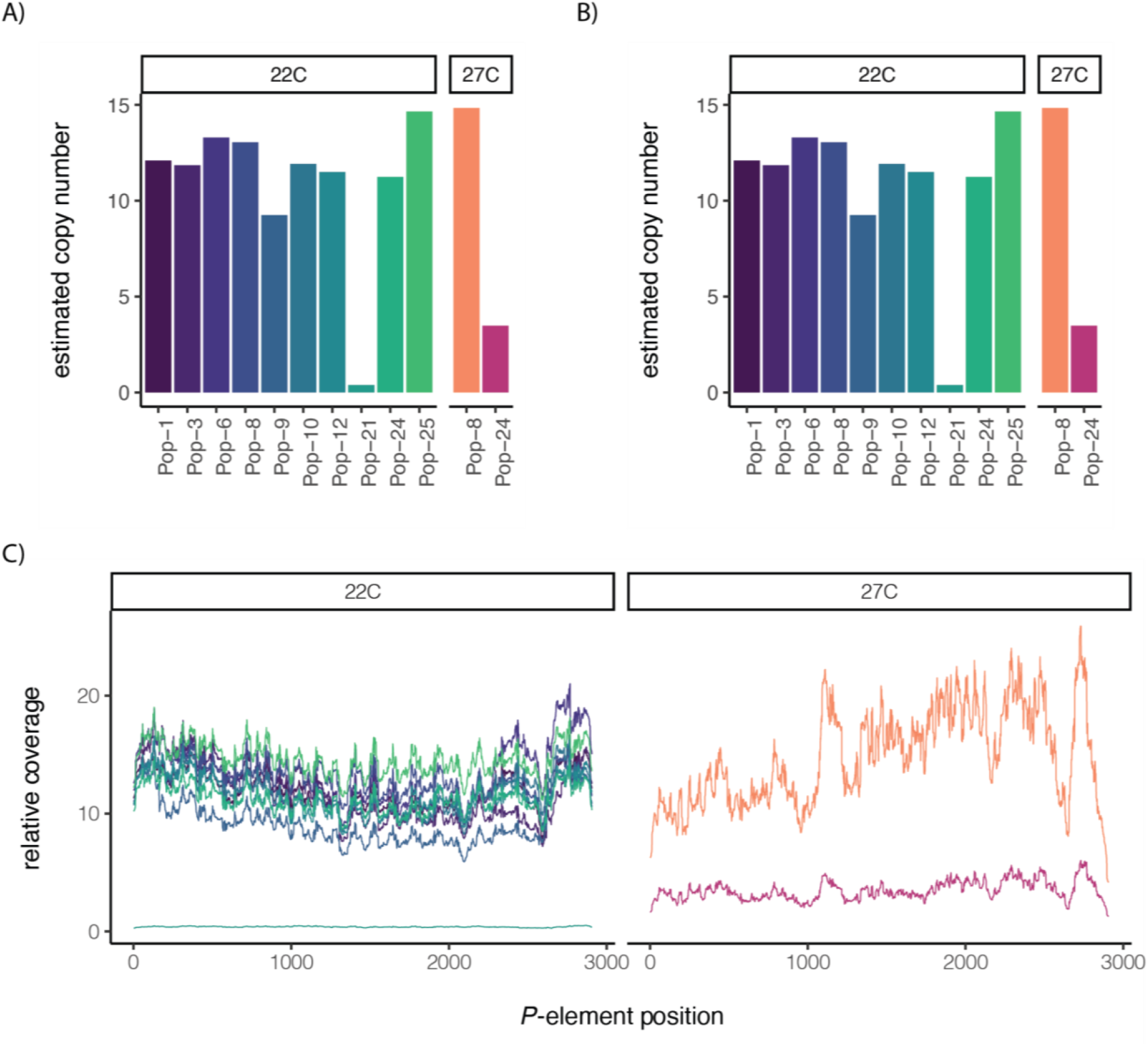
Genomic invasion of *P*-elements in evolved populations. A and B) Estimated copy numbers of genomic P-elements based on A) coverage relative to single copy genes and B) summed frequency of individual P-element insertions. C) Sliding window plot of coverage across the *P*-element consensus sequence.

### Recurrent extinction of colonized populations at 27°C

At 27°C, where *P*-element transposition rates are higher (Moon et al. 2018), evolved induction and repression were not observed with the exception of Population 8 (Figure 1B, Supplementary Table 4). Furthermore, all populations went extinct within 3 to 32 generations of transfer to 27°C (Supplementary Figure 1). When we attempted to re-establish extinct 27°C populations using embryos from the corresponding population at 22°C, extinctions occurred again rapidly, within 4 generations of transfer (Supplementary Figure 1).

To evaluate whether the recurrent extinction of 27°C populations resulted from an overproliferation of *P*-elements we deep sequenced pooled samples (130-174X) of 8 and 9 females, from the final extinction generation of Population 24 (Generation 15) and Population 8 (Generation 41) respectively, yielding 7-10X coverage per haploid genome. We observed that extinct 27°C populations exhibited *P*-element copy numbers that are comparable to (Population 8, G41), or considerably less than (Population 24, G15), populations reared at 22°C (FIgure 2A-B). Furthermore, qPCR of adult genomic DNA from extinction generations at 27°C did not suggest an excess of genomic *P*-elements beyond the copy numbers observed at 22 °C (Supplementary Figure 2). Therefore, the extinction of invaded populations at 27°C does not reflect an excess of accumulated *P*-element copies *per se* (Figure 2C).

It remains possible that the extreme condition represented by 27°C—together with accumulating deleterious *P*-element insertions—drive extinction. Specifically, *P*-element insertions could produce temperature sensitive mutations which are dominant and deleterious at 27°C, but not at 22°C, driving extinction in the former (Suzuki et al. 1967; Holden and Suzuki 1973). This would explain the very rapid extinctions we observed when reestablishing 27°C from 22°C embryos (Supplementary Figure 2). We did not investigate this possibility, but rather focused our analysis on invasion outcomes at 22°C.

### piRNA-mediated repression evolves at 22°C

Increasing maternal repression of hybrid dysgenesis in laboratory populations maintained at 22T suggests the evolution of *P*-element derived piRNA biogenesis. To examine changes in piRNA pools over the course of the experiment, we deep sequenced piRNAs from two populations (1 and 10) at 6 different time points (Figure 3A, Supplementary Table 5, Supplementary Table 6). Both populations exhibited increasing *P*-element derived piRNA production over successive generations. Furthermore, the estimated fraction of *P*-element derived piRNAs that were produced by the ping-pong amplification cycle similarly increased (Figure 3A). piRNA abundance has previously been correlated with ping-pong fraction, and likely reflects increasing efficiency of piRNA production through ping-pong amplification (Kelleher and Barbash 2013).

**Figure 3.**
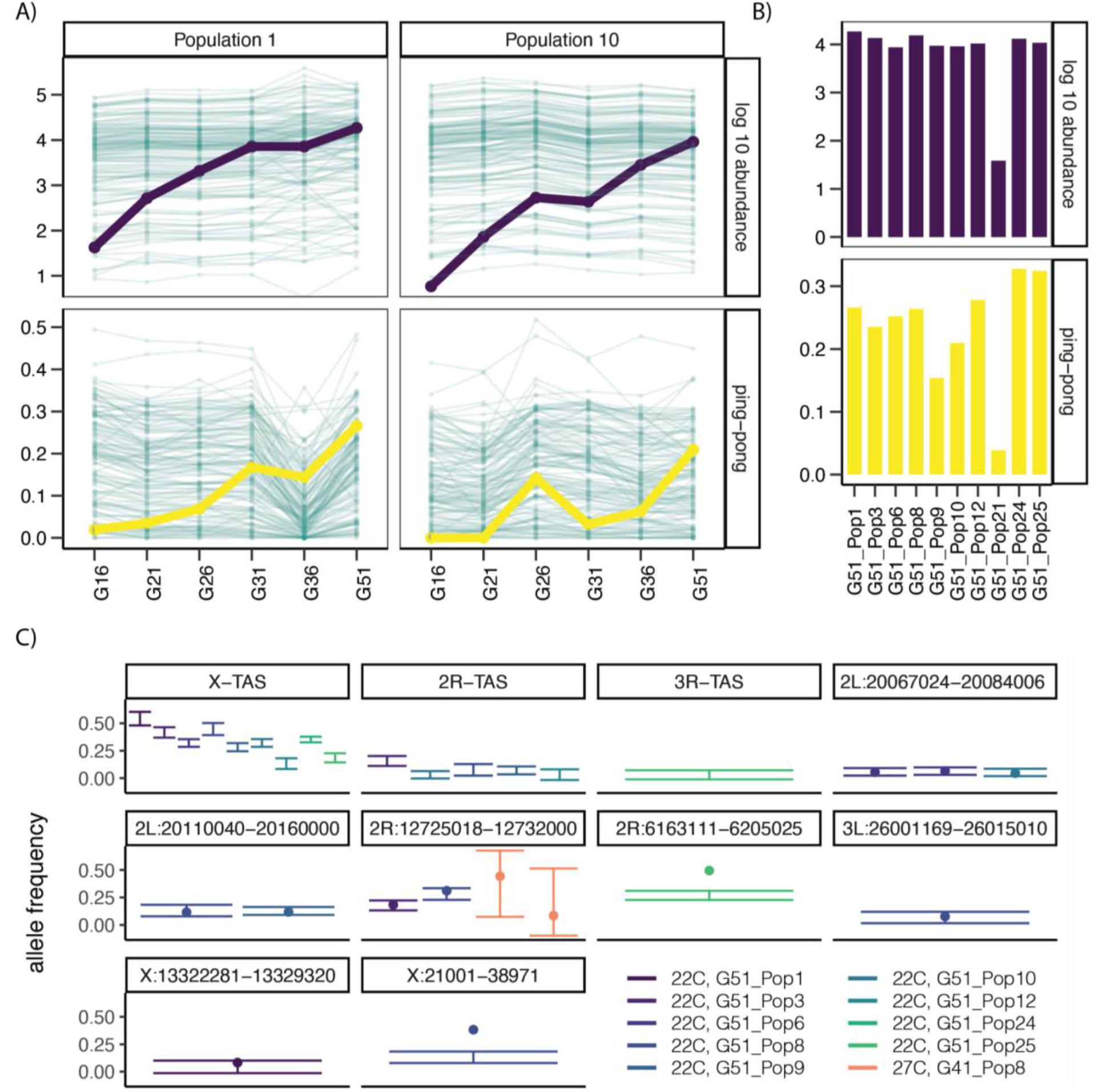
piRNA-mediated silencing in evolved populations. A) Change in *P*-element derived piRNA abundance (purple) and ping-pong fraction (yellow) over successive generations in two populations as compared to all over TE families. Resident (non *P*-element) TE families are represented in the background in green. B) Overall abundance and ping-pong fraction among *P*-element derived piRNAs in 10 populations evolved for 51 generations at 22°C. C) Estimated frequency of insertion alleles into 10 different piRNA clusters in each population. The point indicates the frequency estimated by the proportion of reads overlapping the insertion that support the cluster. The bars indicate a 95% prediction interval for the frequency of the cluster insertion based on a linear model (see methods).

We further connected differences in maternal repression (Figure 1A) with differences in piRNA production and abundance by deep sequencing ovarian piRNAs from all 10 populations at Generation 51. With the exception of population 21, all evolved populations exhibited abundant *P*-element piRNAs with robust ping-pong fractions (Figure 3B, Supplementary Table 5, Supplementary Table 6). In contrast, the population 21 piRNA pool is characterized by reduced *P*-element derived piRNA abundance and ping-pong fraction. Taken together, these data suggest that maternal repression in evolved populations is underpinned by the production of *P*-element derived piRNAs.

Studies in both laboratory (Kofler et al. 2018) and natural populations (Brennecke et al. 2008; Khurana et al. 2011; Zhang et al. 2020) have connected the evolution of *P*-element derived piRNAs with *P*-element insertions into piRNA clusters. In particular, the X telomere associated sequences (*X*-TAS), a subtelomeric satellite array, are an insertion hotspot for *P*-elements (Karpen and Spradling 1992). Furthermore, *P*-element insertions in *X-*TAS isolated from natural populations are known to establish piRNA silencing (Ronsseray et al. 1996; Marin et al. 2000; Stuart et al. 2002). Consistent with these observations, we identified *X-*TAS insertion alleles in all 22°C populations except population 21 (Figure 3C, Supplementary Table 7). Furthermore, we identified *P*-element insertions into *2R* and *3R*-TAS clusters in multiple populations, as well as non-TAS piRNA clusters throughout the genome (Figure 3C). piRNA biogenesis in laboratory populations is therefore associated with insertions into piRNA clusters.

While no *P*-element insertions into piRNA clusters were identified in Population 24 at 27°C, we did identify two insertions into a euchromatic piRNA cluster on chromosome 2R in Population 8 (Figure 3C). The presence of these insertions is consistent with our observation that females from this population exhibited evolved repression (Figure 1B), and may partially explain why Population 8 persisted for longer than others at 27°C.

### *IncRNA:CR4365l* is novel insertion hotspot

In addition to *X-*TAS, *P*-element based transgenes exhibit preferential insertion in portions of the euchromatic genome, with 30-40% of all new insertions occurring in “hotspot” genes (Bellen et al. 2004; Spradling et al. 2011). We observed a similar yet weaker enrichment in our data, with 17% of 4925 insertions we identified in 22°C populations occurring in the top 100 hotspot genes (Figure 4A, Supplementary Table 9). Similarly, we observed that genomic windows containing previously identified hotspots and hotpoints (preferred insertion nucleotide positions) are greatly enriched for *P*-element insertions in 22°C populations. 5.5% of genomic windows overlapping hotspots or hotpoints contain a *P*-element insertion allele in at least one population, as compared to 1.5% of windows that do not overlap preferred sites (*X*^2^ = 681.08, *df* = 1, *P*-value < 10^-15^, Figure 4A).

**Figure 4.**
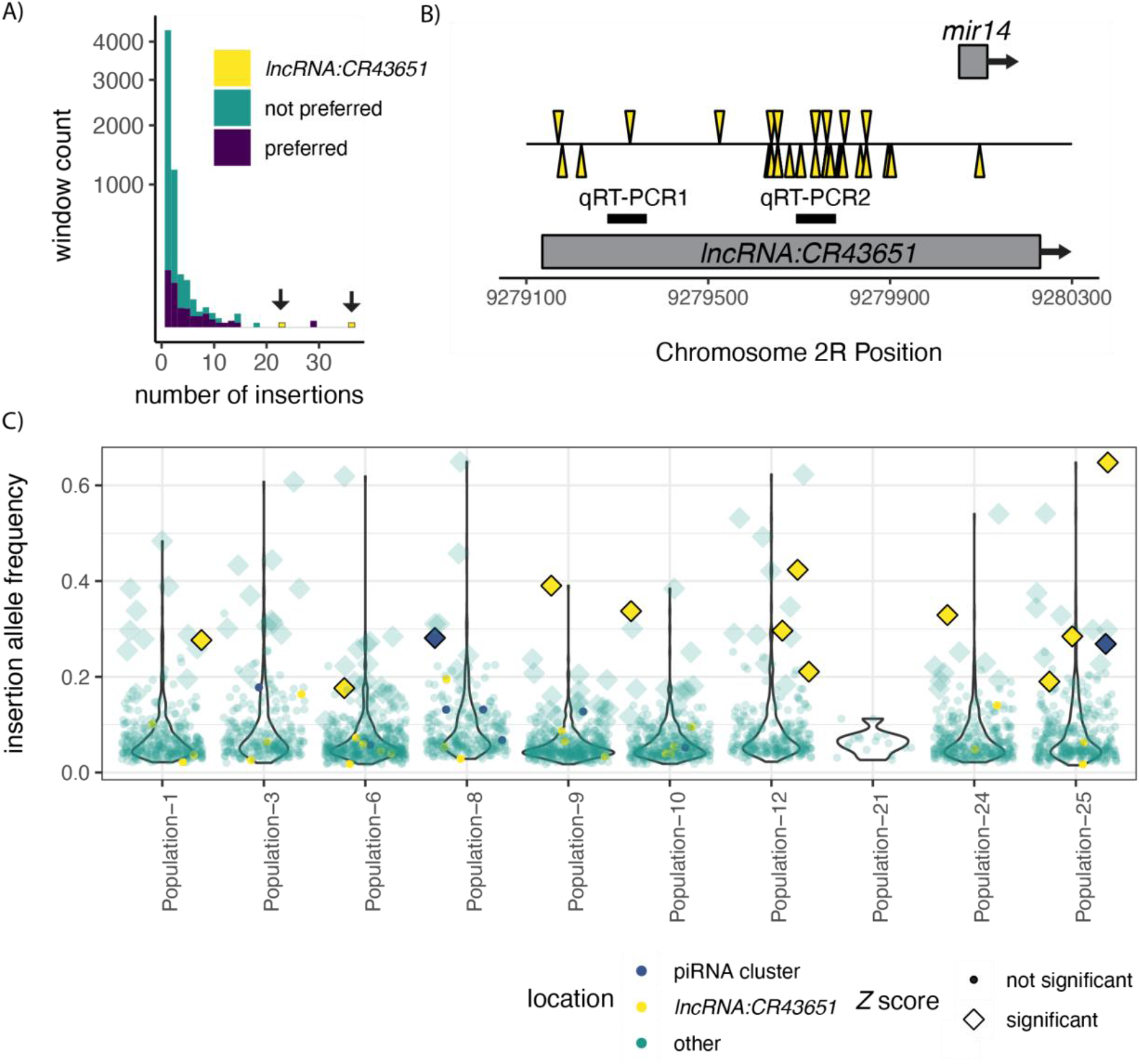
*lncRNA:CR43651* is an insertion spot and contains multiple high frequency insertion alleles. A) Frequency of 1 kb genomic windows containing different numbers of P-element insertion alleles in the 22°C population at generation 51. Preferred genomic windows overlap previously identified insertion hotspots and hotpoints (Spradling et al. 2011). Arrows point to two genomic windows containing regions overlapping *lncRNA:CR43651*, that contain an unusually high number of insertions. B) *P*-element insertion alleles upstream and within *mir-14* are indicated. Plus strand insertions, transcribed in the same direction as *mir-14* and *lncRNA:CR43651* are indicated above the line, while minus strand insertions are indicated below the line. Black lines indicate qPCR products quantified in Figure 5I. C) The frequencies of individual insertion alleles in *lncRNA:CR43651*, piRNA clusters, and elsewhere in the genome are indicated for each population. Insertion alleles with unusually high frequencies (*Z* score > 1.96, *P*-value < 0.05) are indicated.

The genomic window that is most densely populated with *P*-element insertions in our 22°C populations occurs upstream of *mir-14* and within the transcript body of *lncRNA:CR43651* (Figure 4B). This 1 kb region contains a total of 36 insertion alleles, with every population except population 21 segregating 3 or more alleles (Figure 4C). *lncRNA:CR43651* is not among the 100 most enriched hotspots identified in the *D. melanogaster* genome (Spradling et al. 2011). However, the sheer number of insertions, including insertions that occur on opposing strands in the same position (Supplementary Table 9), suggests that this region was a preferred insertion site in our experiment.

### Insertions in *lncRNA:CR43651*, but not piRNA clusters, are targets of selection in laboratory populations

While an insertion bias may partially explain the occurrence of *P*-elements in *lncRNA:CR43651*, selection may also have acted to retain those alleles and increase their frequency in laboratory populations. We therefore considered whether insertion alleles in this region might segregate at unusually high frequencies, when compared *P*-element insertions elsewhere in the genome from the same population. To this end, we estimated the allele frequency of every *P*-element insertion allele in each population based on 1) the proportion of reads overlapping the insertion site that support the insertion allele and 2) a regression model of allele frequencies onto raw read support for each insertion (materials and methods). We then calculated a *Z*-score for the estimated frequency of each *P*-element insertion allele (Supplementary Table 9). A high *Z*-score indicates a *P*-element that is segregating at exceptionally high-frequency, a potential indicator of positive selection.

The two approaches for estimating *P*-element insertion allele frequency were highly concordant, and identified 113 insertion alleles (out of 2901) whose estimated allele frequency was unusually high (*Z*-score > 1.96, *P-*value < 0.05) for both allele frequency estimates. The proportion of reads method identified one additional high frequency insertion, while the regression based estimate identified 306 additional high-frequency insertion alleles. Of the 113 insertion alleles where both frequency estimates were high-frequency outliers, 11 correspond to insertion alleles in *lncRNA:CR43651* (Figure 4C). This represents a highly significant enrichment of high frequency alleles among *lncRNA:CR43651* insertions (*X*^2^ = 6.18, *df* = 1, *P-value* < 10^-15^).

We next considered whether the population frequency of *P*-element insertion alleles in *lncRNA:CR43651* might be elevated by an insertion bias. Recurrent insertion in the same “hotpoint” nucleotide position can potentially increase allele frequencies independently of selection (Zhang et al. 2020). However, in our laboratory populations insertion alleles in previously identified hotpoint sites do not exhibit higher allele frequencies than those outside of hotpoint sites (two-sample permutation test, all populations, P > 0.05). Therefore, the unusually high frequency of *lncRNA:CR43651* insertion alleles cannot be explained by recurrent insertion into the same nucleotide position, and rather suggest these alleles are targets of positive selection.

Simulation models predict that TE insertions into piRNA clusters are also targets of weak positive selection if they reduce the occurrence of dominant deleterious mutations or hybrid dysgenesis (Kelleher et al. 2018; Kofler 2019). We therefore examined whether any *P*-element insertions in piRNA clusters represented high frequency outliers (Figure 3C). Insertion alleles into *X, 2R* and *3R*-TAS were excluded from this analysis, since we are unable to resolve insertions that occur in different satellite repeat units. Of 11 remaining non-TAS piRNA cluster insertion alleles, 2 corresponded to allele frequency outliers, a marginally significant enrichment (Fisher’s Exact Test, *P*-value = 0.065, Figure 4C). Therefore, while insertion alleles in piRNA clusters may experience weak positive selection in some populations, our observations suggest that selection on insertion alleles in *lncRNA:CR43651* is considerably stronger.

### Hotspot insertions are associated with piRNA biogenesis from *lncRNA:CR43651*

Signatures of selection on insertions in *lncRNA:CR43651* implies they have a beneficial functional effect on a proximal gene product. Transposon insertions can alter gene function through disrupting functional sequences, introducing new cis-regulatory sequences or exons, and initiating the formation of heterochromatin (reviewed in Feschotte 2008). Given the location of high frequency insertions within *lncRNA:CR43651* and upstream of *mir-14, P*-element insertions could impact the transcription or function of one or both of these transcripts. Both *mir-14* and *lnRNA:CR43651* are broadly expressed in many larval and adult tissues, including the adult ovary (Supplementary Figure 3). We therefore took advantage of our ovarian RNA samples and small RNA data to evaluate whether expression of *mir-14* or *lncRNA:CR43651* is altered by *P*-element insertions. If *P*-element insertions impact the expression of either transcript, we predict 1) altered expression in populations that have multiple high frequency insertions and 2) a successive change in expression in those same populations over the course of the experiment.

We did not uncover any evidence that *P*-element insertions impact the expression of *mir-14* in our small RNA data. While both the 5-prime (*mir-14-RB*) and 3-prime (*mir-14-RA*) species of *mir-14* varied in expression between populations, there is no obvious relationship between expression level and the presence or allele frequency of *P*-element insertions in this region. For example, populations 12 and 25 both have multiple high frequency *P*-element insertions at Generation 51 (Figure 4C), but highly dissimilar expression levels of *mir14-RB* (Figure 5A). Furthermore, variation in *mir-14* expression levels over successive generations in population 1 and 10 show no particular trajectory of increase or decrease (Figure 5B). Nevertheless, because our small RNA data were from adult ovaries, we cannot rule out impacts in a subset of germline cells or in other adult or larval tissues.

**Figure 5.**
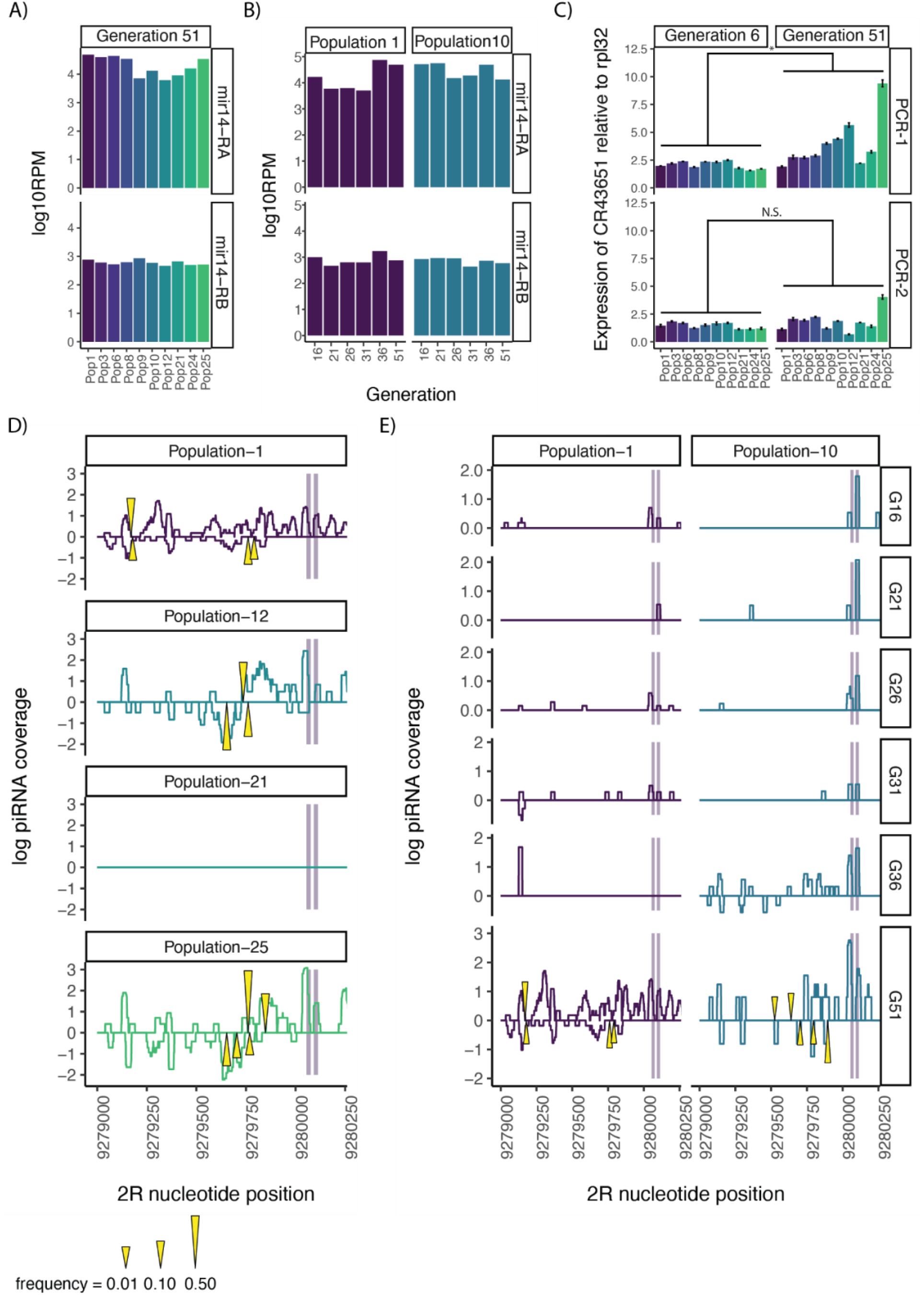
Transcriptional impacts of *P*-element insertions in *lncRNA:CR43651*. A-B) Relative abundance of the 3-prime (*mir14-RA*) and 5-prime (*mir14-RB*) miRNA products of *mir14* from all populations at Generation 51 (A) and successive successive generations in populations 1 and 10 (B). C) qRT-PCR of two RNA fragments from *lncRNA:CR43651* represented in Figure 4B in pooled ovarian samples from generation 6 and generation 51 for all populations. D-E) piRNAs from both genomic strands of a portion of *lncRNA:CR43651* represented in Figure 4B. Locations of *mir-14* transcripts are highlighted in grey. Locations of *P*-element insertions in each population in generation G51 are indicated by yellow triangles, which are scaled according to the estimated insertion frequency. Abundance is in reads per million mapped miRNAs. piRNA data are from representative populations at Generation 51 (D) or over successive generations in populations 1 and 10 (E).

In contrast to *mir-14*, the expression and structure of the *lncRNA:43651* does appear to be impacted by *P*-element insertions. RNA fragments from a region upstream of most *P*-element insertions (qRT-PCR1) show increased abundance in ovarian RNA samples from Generation 51 as compared to Generation 6, with particularly pronounced increases occurring in Populations 12 and 25 (*F1,9* = 6.75, *P* = 0.03, Figure 5C). This surprisingly suggests that *P*-element insertions in *lncRNA:CR43651* increase the expression or stability of transcripts from adjacent sequence. In contrast, RNA fragments from a region containing numerous *P*-element insertion sites (qRT-PCR2) show no systematic change in expression: with individual populations showing increased and decreased expression in Generation 51 (*F1,9* = 1.28, *P* = 0.29, Figure 5C). The latter observation likely reflects the disruption of the qRT-PCR2 assay by the occurrence of *P*-element insertions at primer target sites and within the amplicon.

Increased abundance of *lncRNA:CR43651* fragments could reflect the transcription of precursor piRNAs from within silenced *P*-element insertions. piRNA mediated silencing of euchromatic TEs often results in the formation of *de novo* piRNA clusters, which exhibit bi-directional transcription of precursor piRNAs that extends into adjacent sequences (Shpiz et al. 2014). Consistent with this model, we observed piRNA biogenesis from the *lncRNA:CR43651* region surrounding *P*-element insertions in all populations except population 21, suggesting that piRNA biogenesis emerges concomitantly with *P*-element insertion (Supplementary Figure 4, Figure 5D). Similarly, piRNA biogenesis is largely confined to the region surrounding *mir-14* in Populations 1 and 10, but throughout the *P*-element insertion region of *lncRNA:CR43651* increases over successive generations (Figure 5E). Finally, we detected *lncRNA:CR43651-P*-element chimeric transcripts in cDNA samples from populations 1, 12 and 25 at generation 51, but not generation 6, linking piRNA biogenesis to the production of putative precursors including both TE and *lncRNA:CR43651* sequences (Supplementary Figure 5). As expected, no chimeric transcripts were detected in cDNA from population 21, which lacked *P*-element insertions in *lncRNA:CR43651* and associated piRNA biogenesisx. Collectively, these observations support the production of precursor piRNAs including *lncRNA:CR43651* fragments as a result of *P*–element insertions.

## Discussion

Through our three year laboratory evolution experiment we uncovered several key findings that advance and refine our understanding of the genomic and phenotypic consequences of a novel TE invasion. We observed that the invading TE rapidly expanded in genomic copy number (Figure 2B), ultimately leading to both sterility effects (Figure 1B) and its own repression through insertions into piRNA producing sites (Figure 3). These observations are consistent with previous laboratory evolution experiments (Kidwell et al. 1988; Kofler et al. 2018), stochastic simulation models (Kelleher et al. 2018; Kofler 2019), and surveys of natural variation following *P*-element invasion (Hill et al. 2016; Zhang et al. 2020), all of which demonstrate that the emergence of dysgenic phenotypes are quickly followed by evolved piRNA mediated repression. However, in this study we also made the novel and surprising observation that *P*-element invasion can produce clusters of beneficial insertions at the same locus, resulting in parallel adaptation in replicate populations on a very short evolutionary time scale. These observations raise exciting new questions about how the TE invasion events play in shaping the landscape of molecular adaptation.

Previous studies of *P*-element invasion focus on the evolution of repression of the invading TE. A critical and contentious question in the evolution of host TE repression is the degree to which repressor alleles are targets of natural selection (Charlesworth and Langley 1986). We uncovered numerous *P*-element insertions into piRNA clusters in our experimental populations, which represent repressor alleles. However, few of them were segregating at high frequencies indicative of natural selection. Overall therefore, our results support the model in which while repressor alleles are weakly beneficial, the evolution of TE repression is largely mutation driven, based on the high rate of insertion of TEs into piRNA producing sites (Charlesworth and Langley 1986; Kelleher et al. 2018; Kofler 2019; Zhang et al. 2020). These results furthermore are consistent with our previous analysis of *P*-element insertions in piRNA clusters in North American *D. melanogaster*, which uncovered numerous insertion alleles which appear to be neutral (Zhang et al. 2020).

Here we revealed that the exceptional transposition rates that follow TE invasion not only drive the evolution of repression, they also provide fuel for other forms of adaptation. Specifically, we discovered 36 insertion alleles into *lncRNA:CR43651* in 9 separate populations, of which 11 were high frequency outliers (Figure 4). The recurrence of these insertions into the same small genomic window (~1 kb, Figure 4B) likely in part reflects the known insertion preference of *P*-elements for origins of replication (Spradling et al. 2011), since *lncRNA:CR43651* occurs in an origin of replication (Supplementary Figure 6). Nevertheless, this region is not among the top 100 *P*-element insertion hotspots from transgenic screens (Bellen et al. 2004, 2011; Spradling et al. 2011), arguing that insertion preference alone is not sufficient to explain the exceptional number of insertion alleles in our experiment (Figure 4A,4B). Rather insertion bias, together with selection for beneficial insertion alleles (Figure 4C) must have occurred. This synergism between insertion site preference and occurrence of beneficial insertions echoes a series of *roo* solo-LTR insertions surrounding *CG18446*, one of which exhibits signature of positive selection and impacts cold tolerance by increasing *CG18446* expression (Merenciano et al. 2016).

Why are *P*-element insertions into *lncRNA:CR43651* likely beneficial in laboratory populations undergoing *P*-element invasion? An appealing explanation is that the insertions compensate for the deleterious fitness effects of *P*-element transposition on fertility by upregulating *mir-14*, a known repressor of apoptotic cell death (Xu et al. 2003). However, insertions with such an effect are also predicted to be beneficial in natural populations that exhibited hybrid dysgenesis following *P*-element invasion, such as North America (Kidwell 1983; Kidwell, M.G., Frydryck, T., and Novy, J.B. 1983). This is not the case: as only one of >200 fully sequenced North American lines contains an insertion in this gene region (Mackay et al. 2012; Zhang et al. 2020). The absence of evidence for selection on *lncRNA:CR43651* insertions in natural populations would argue that they confer an advantage specifically under laboratory conditions, but not in nature.

An alternative explanation, for which we have no direct evidence, is that *P*-element insertions into *lncRNA:CR43651* impacts a trait that would be strongly selected in a lab selection experiment such as development time or fecundity. Interestingly, *mir-14* is also a negative regulator of Ecdysone Receptor in *D. melanogaste*r (Varghese and Cohen 2007), which regulates several key physiological processes including pupation and oogenesis (reviewed in Thummel 1995; Schwedes and Carney 2012; Finger et al. 2021). If *P*-element insertions exert their beneficial regulatory effects in non-ovarian tissue, such as in whole larvae, this would explain the absence of a systematic effect on *mir14* expression in the ovary (Figure 5A, 5B).

In summary, our work demonstrates that TE invasions not only produce abundant *de novo* insertions that establish piRNA-mediated repression, they may also drive the production of novel insertion alleles that fuel adaptation. While there is extensive work on adaptive TEs insertions in natural populations (reviewed in González and Petrov 2009; Casacuberta and González 2013), little is known about how insertion biases and genome invasions shape the landscape of these adaptive insertions. Our work here shows that genome invasions can accelerate TE-driven adaptation to a rate that vastly outpaces that of nucleotide substitutions and structural mutations. The degree to which TE-dependent adaptation coincides with genome invasions represents an interesting direction for future work.

## Materials and Methods

### Establishment and maintenance of populations with *P*-element invasion

We used a highly inbred version of a *P*-element free wild type strain (KSA, Bloomington #3862), which was generously provided by Tony Long. *P*-elements were introduced into the KSA using germline transformation with a wild-type full-length *P*-element clone (pπ25.1; O’Hare and Rubin 1983) in a pUC13 plasmid, which was generously provided by Michael Simmons. The plasmid was injected at a concentration of 300 ng/μL by BestGene Inc (Chino Hills, CA). Male and female offspring of injected embryos were crossed to uninjected A4, and resulting stocks were screened for *P*-elements by PCR. Ten independent transformations were detected in *P*-element positive offspring of separate embryos, and were used to establish 10 replicate populations. Each population was expanded at room temperature for 3 generations, and then split into two replicates, which were maintained at low (22°C) and high temperature (27°C).

Every generation, ~1000 eggs were collected from each population to ensure large and consistent population sizes across replicate populations and treatments. Two days before egg collection, newly-eclosed (1-4 day old) adults were placed in population cages and provided with grape juice agar plates supplemented with yeast paste, to promote oogenesis. One day before egg collection, additional plates and yeast paste were provided, and the females were allowed to lay for 24 hours. Eggs were collected by flushing the yeast paste along with the eggs into a collection container with a filter net. Collected eggs were rinsed and 200 uL of eggs were transferred into bottles, which we determined corresponded to about 250 embryos. 4 replicate bottles were established on standard cornmeal media, which after eclosion represented approximately 1000 adult flies to begin the next experimental generation. Flies were maintained for 51 generations.

### Phenotypic measurements

Every 5 generations, we measured the induction of hybrid dysgenesis associated with the *P*-element for each population at each temperature by evaluating the ability of experimental males to cause dysgenesis among their F1 offspring females when crossed to M strain females (Canton-S). We similarly measured repression by evaluating the ability of experimental females to repress hybrid dysgenesis in their F1 offspring females when crossed to P strain males (Harwich). To capture variation in induction and repression within each population: 20 replicate crosses including an individual experimental male or female and 1-2 mating partners (Canton-S females/Harwich males) were performed. Crosses were maintained at 29°C, and the proportion of F1 ovarian atrophy was estimated from a sample of 20 female offspring.

The change of repression or induction over generations was modeled with a mixed effects logistic regression model in the R package lme4 (Bates et al. 2015). The model was fit separately for each population and cross direction follows:

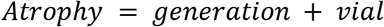

Generation was treated as a fixed effect and vial as a random effect.

### *P*-element copy number estimation

Pooled samples of genomic DNA from extinct populations reared at 27°C were used as templates for quantitative PCR. Abundance of *P*-element and *rpl32* in genomic DNA was estimated using SYBR green PCR mastermix (Applied Biosystems) according to manufacturer instructions. Three technical measurements were taken for each sample, and relative abundance of *P*-elements as compared to *rpl32* was estimated based on a five-point standard curve. Primers were as follows: *P*-element-F: 5’-CACCGAAATGGATGAGTTGACG-3’, *P*-element-R: 5’-TAATAAGTCCGCCGTGAGACAC-3’, *rpl32-F:* CCGCTTCAAGGGACAGTATC, and *rpl32*-F: GACAATCTCCTTGCGCTTCT.

To convert *P*-element abundance into genomic copies, we also generated a *P*-element abundance estimate for females from the *TP5, w1* (BDSC 64168), a stock homozygous for a single non-autonomous P-element in the *X*-TAS. An estimate of the average *P*-element copy number in each population was obtained by dividing its normalized abundance by that of TP5 and multiplying by 2.

### Whole genome sequencing

Pooled samples of genomic DNA from experimental populations were to construct genomic sequencing libraries by using the Nextera DNA Flex Library Preparation Kit according to manufacturer instruction (Illumina). Pooled samples from the last generation before extinction of two 27°C populations (9 females, population 8; 8 females, population 24), and from generation 51 of all the 22°C populations (20 females/population) were examined. DNA was extracted from whole flies using QIAGEN DNeasy Blood & Tissue Kits and used as a template for library construction. Adapter-tagged DNAs were washed and amplified by PCR. Final libraries were sent for whole genome sequencing on an NextSeq 500 (27°C populations, High Output, Paired-End 75) and Illumina NovaSeq 6000 (22°C populations, S4, Paired End 150) by Genomic and RNA Profiling Core in Baylor College of Medicine.

### Detecting *P*-element insertions

*P*-element insertions were detected as described in (Zhang and Kelleher 2017; Zhang et al. 2020). Whole genome sequencing reads were locally aligned to full-length *P*-element consensus sequence (O’Hare and Rubin 1983) using Bowtie 2 (Langmead and Salzberg 2012) to identify reads that include *P*-element sequence. *P*-element sequences were then trimmed from aligned reads. The remaining sequences were aligned to the A4 assembly (GCA_003401745.1, Chakraborty et al. 2019) using Bowtie2 to identify insertion sites. Telomeric-associated sequences (TAS), which are repetitive satellites containing preferred insertion sites for *P*-elements (Karpen and Spradling 1992), were masked from the assembly (X-TAS, CM010541.1:29332..35458; 2L-TAS, CM010542.1:38964..39347). Alignments with MAPQ>20 and edit distance <4 were retained as used to identify *P*-element insertion sites using PIDFE (Zhang and Kelleher 2017; Zhang et al. 2020).

*P*-element reads that did not align to the masked A4 assembly were next aligned to a custom library of TAS sequences using Bowtie 2 and outputting all valid alignments (-a). The custom library was comprised of TAS regions from the A4 assembly (X-TAS, CM010541.1:29332..35458; 2L-TAS, CM010542.1:38964..39347), or the dm6 reference genome (2R-TAS, 2R: 25258060..25261551; 3R-TAS, 3R: 32073015..32079331; 3L-TAS, 3L: 1..19608; Walter et al. 1995; Yin and Lin 2007; Hoskins et al. 2015). A read was considered to support a TAS insertion if aligned to any TAS sequence with an edit distance was fewer than 4.

### Estimating the frequency of insertion alleles

To estimate the frequency of *P*-element insertion alleles in our pooled population samples, we first generated pseugenomes for each annotated *P*-element insertion in a given population using make_P_pseudo_only.pl from the PIDFE package (Zhang and Kelleher 2017; Zhang et al. 2020). Each pseudogenome included the *P*-element itself, an 8 nucleotide target site duplication, and 1000 nt of flanking genomic sequence from the A4 assembly. Reads were aligned concordantly to a combined library containing the unmasked A4 assembly and pseudogenomes for all insertions using Bowtie 2. Read pairs aligning to an insertion pseudogenome and overlapping the insertion site with an edit distance <4 and a quality score >20 were retained and considered to support the insertion.

Next, we identified read pairs supporting the un-inserted reference allele by aligning read pairs that did not support an insertion allele concordantly to the A4 reference genome only, using Bowtie 2. Read pairs aligning to the reference genome and overlapping the site with an edit distance <4 and a quality score of >20 were considered to support the reference allele.

Based on the number of read pairs supporting the insertion allele and the reference allele, we estimated allele frequencies in two ways: a proportion of reads approach and a regression approach. For the proportion of reads approach the number of reads supporting the insertion allele was divided by the total number of reads supporting the insertion or reference allele for each insertion position using the script newfreq.pl, and updated version of a frequency calculating script from PIDFE that runs faster by acting as a wrapper for samtools. For insertion sites represented by two insertion alleles (sense and antisense strand), the frequency was subsequently recalculated to account for reads corresponding to the alternative insertion allele (e.g. freq = insertion allele sense / (insertion allele sense + insertion allele antisense + reference allele).

To obtain a second estimate of the frequency of each insertion allele, as well as to identify alleles whose frequency may have been over or underestimated, we further pursued a regression approach. First we fit a linear model of insertion allele frequency as a function of the read count supporting the insertion allele separately for each population in R. Based on the regression line, a predicted allele frequency was obtained based on the read count supporting each insertion. Furthermore, insertion alleles whose estimated frequency was outside of the 95% prediction interval for a given population were identified as outliers, whose frequency was likely incorrectly estimated by calculating the fraction of reads supporting the reference allele.

### Sliding window coverage analysis

To generate sliding window coverage plots for *P*-elements in each population, Bedtools genomecov was used to calculate coverage at each position, based on alignments of individual unpaired reads to the *P*-element consensus (-d; Quinlan and Hall 2010). Coverage was scaled to the total number of aligned reads in each pooled genome resequencing library to generate a reads per million mapped coverage estimate (-scale 10^-6^/total alignable reads).

### Analysis of allele frequencies

We evaluated the relationship between insertion allele frequencies (proportion of reads method only) and recombination rate by taking advantage of the high resolution recombination made from Comeron *et al*. (2012). dm3 coordinates with known recombination rates were transferred first to the dm6 assembly using the flybase coordinates converter tool. Windows were subsequently translated to A4 using the NCBI genome remapping tool Remap. Annotated insertions in each population were then associated with the local recombination rate using the R package GenomicFeatures (Lawrence et al. 2013). The relationship between insertion allele frequency and local recombination rate was evaluated through an extra sum-of-squares *F*-test comparing a null model containing only an intercept to an alternative model in which the frequency is a function of recombination rate. Insertions with unusual allele frequencies (outside the 95% prediction interval of the linear model) were excluded from this analysis.

To evaluate the relationship between insertion allele frequency and previously identified insertion hotspots and hotpoints, we first translated hotspot/hotpoint locations into A4 coordinates using NCBI Remap. We then identified insertions located at hotpoints or in hotspots using GenomicFeatures. For hotpoints, we considered a 20 nt window surrounding each hotpoint position to account for potential uncertainties about the precise hotpoint position in A4. The relationship between insertion allele frequency and presence in a preferred insertion site (hotspot or hotpoint) was evaluated using a permutation test in R (oneway_test, coin package, Zeileis et al. 2008).

To identify insertions with unusually high frequencies, we calculated *Z*-scores for the log transformed frequencies of all insertion alleles based on the mean and variance in each population. Alleles with positive *Z*-scores and low *P-*values were considered unusually high frequency. This analysis was performed separately for both methods of estimating allele frequency. The occurrence of all insertion alleles in piRNA clusters and within *lncRNA:CR43651* was determined using GenomicFeatures.

### Small RNA sequencing

Ovaries from: 1) generation 51 females of all the 22°C populations and 2) generations 6, 21, 26, 31, and 36 of populations 1 and 10, were used to build small RNA libraries. 20 3-6-day old female ovaries were dissected from each population, placed directly in Trizol reagent (Invitrogen), and homogenized. Small RNA libraries were prepared as described in (Wickersheim and Blumenstiel 2013). In brief, total RNAs were extracted according to the manufacturer’s instructions, and size fractionated on a 12% polyacrylamide/urea gel to select for 18–30 nt small RNAs. Small RNAs were treated with 2S Block oligo (5’-TAC AAC CCT CAA CCA TAT GTA GTC CAA GCA/3SpC3/-3’), and were subsequently ligated to 3’ and 5’ adaptors, reverse transcribed and PCR amplified using NEBNext Multiplex Small RNA Library Prep Set for Illumina. Small RNA libraries were further purified from a 2% agarose gel and sequenced on a Illumina NextSeq 500 at the University of Houston Seq-N-Edit Core.

### Small RNA data analysis

3’ Illumina adaptors (AGATCGGAAGAGCACACGTCT) were removed from sequencing reads by Cutadapt (Martin 2011). Sequence alignments were made by Bowtie (Langmead et al. 2009). Contaminating ribosomal RNAs were identified and removed by mapping sequencing reads to annotated ribosomal RNAs (dm6) from flybase (Gramates et al. 2017). TE-derived piRNAs were identified by aligning sequencing reads ranging from 23–30 nucleotides (nt) to Repbase (Bao et al. 2015), allowing for up to 2 mismatches. The number of reads mapped to each TE family were counted using a Linux shell script. Redundant TE families in Repbase were identified by checking sequence identity (those consensus sequences that were >90% identical across >90% of their length were categorized as the same TE family). Reads mapped to multiple redundant TE families were counted only once. Reads mapped to multiple non-redundant TE families were discarded. To identify miRNAs, sequencing reads between 18 and 22 nt were aligned to a miRNA reference sequence (dm6) from Flybase (Gramates et al. 2017). TE families or genes with low read count (< 50 on average) in every library were discarded. piRNA counts for each TE family were normalized to the total number of sequenced miRNAs from each library. Normalized read counts were used for comparisons of the abundance of piRNAs between libraries.

Ping-pong fraction was calculated as described in (Brennecke et al. 2008). In brief, small RNA sequencing reads ranging from 23–30 nt were aligned to TE consensus sequences from Repbase (Bao et al. 2015), and redundant TE families in Repbase were identified as described above. For each piRNA, the proportion of overlapping antisense binding partners whose 5’ end occurs on the 10th nucleotide was determined. This fraction was subsequently summed across all piRNAs from a given TE family, while incorporating the difference in sampling frequency between individual piRNAs. Finally, this sum was divided by the total number of piRNAs aligned to the TE family of interest. For multi-mappers, reads were apportioned by the number of times they can be aligned to the reference.

We used the proTRAC workflow to identify piRNA clusters in each population and generation (Rosenkranz and Zischler 2012). Briefly, putative piRNAs were first filtered for redundant sequences (TBr2_collapse.pl) and low-complexity reads (TBr2_duster.pl). Remaining reads were then aligned to the A4 assembly (GCA_003401745.1; Chakraborty et al. 2019) allowing for 0 mismatches using Bowtie (-v 0; Langmead et al. 2009). Alignments were analyzed by proTRAC (version 2.3.1) to identify piRNA clusters, allowing up to 5% of the genome to be annotated as a piRNA cluster (pdens = 0.05), and without considering strand bias (-clstrand 0.5).

### *lncRNA:CR43651* transcripts

Aliquots of the pooled ovarian RNA samples used for small RNA library prep were also used to detect and quantify transcripts of *lncRNA:CR43651* using RT-PCR and qRT-PCR. Briefly, >2 μg total ovarian RNA was reverse transcribed using superscript IV (Invitrogen) using random primers and according to manufacturer instructions. cDNA samples were treated with RNAseH and diluted in water. Abundance of *lncRNA:CR43651* and *rpl32* transcript was estimated using SYBR green PCR mastermix (Applied Biosystems) according to manufacturer instructions. Three technical measurements were obtained from 1:400 diluted cDNA for each population and generation according to a five-point standard curve (1:8, 1:40, 1:200, 1:1000, 1:5000). Primers were as follows: *lncRNA:CR43651-F1:* 5’-GAGAGAGAGAGCGCGTCCTA-3’, *lncRNA:CR43651-R1:* 5’-GGCAAAGTTCACTGGGAAAA-3’, *lncRNA:CR43651-*F2: 5’-TTCACCCAAATCCAAAGAGAAT-3’, *lncRNA:CR43651*-R2: 5’-GTGCGTAGGCATGTATGAGTGT-3’, *rpl32-F:* CCGCTTCAAGGGACAGTATC, and *rpl32*-F: GACAATCTCCTTGCGCTTCT.

The relationship between population, generation and normalized *lncRNA:CR43651* was explored for each qRT-PCR reaction separately using linear regression. Briefly, a linear model containing population as the only explanatory variable was compared to a model containing population and generation. Linear models were fit using lm, and compared using an extra-*SS F*-test in R 4.1.1 (R Core Team 2021)

To detect chimeric transcripts containing both *lncRNA:CR43651* and *P*-element sequence, we performed standard PCR on reverse transcribed cDNA. Four PCR assays were performed, which combined *lncRNA:CR43651-F1* with 4 different primers located close to the 5’ and 3’ ends of the *P*-element sequence: *P*-element-5F: 5’-TTGCTGCAAAGCTGTGACTG-3’, *P*-element-5R: 5’-AGTTTTCACCAAGGCTGCAC-3’, *P*-element-3F: 5’-TGATGAGCCTGTCGATGATGAG-3’, *P*-element-3R: 5’-ATCGCATCCTCCGTCAACTC-3’. Bright bands from the *P*-element 5R reaction indicated the production of chimeric transcripts including sense orientation *P*-element insertions. Bright bands from the *P*-element 3F reaction indicated the production of chimeric transcripts including antisense orientation *P*-element insertions.

