## Supplementary Figure 3 for "TE invasion fuels molecular adaptation in laboratory populations of *Drosophila melanogaster*"

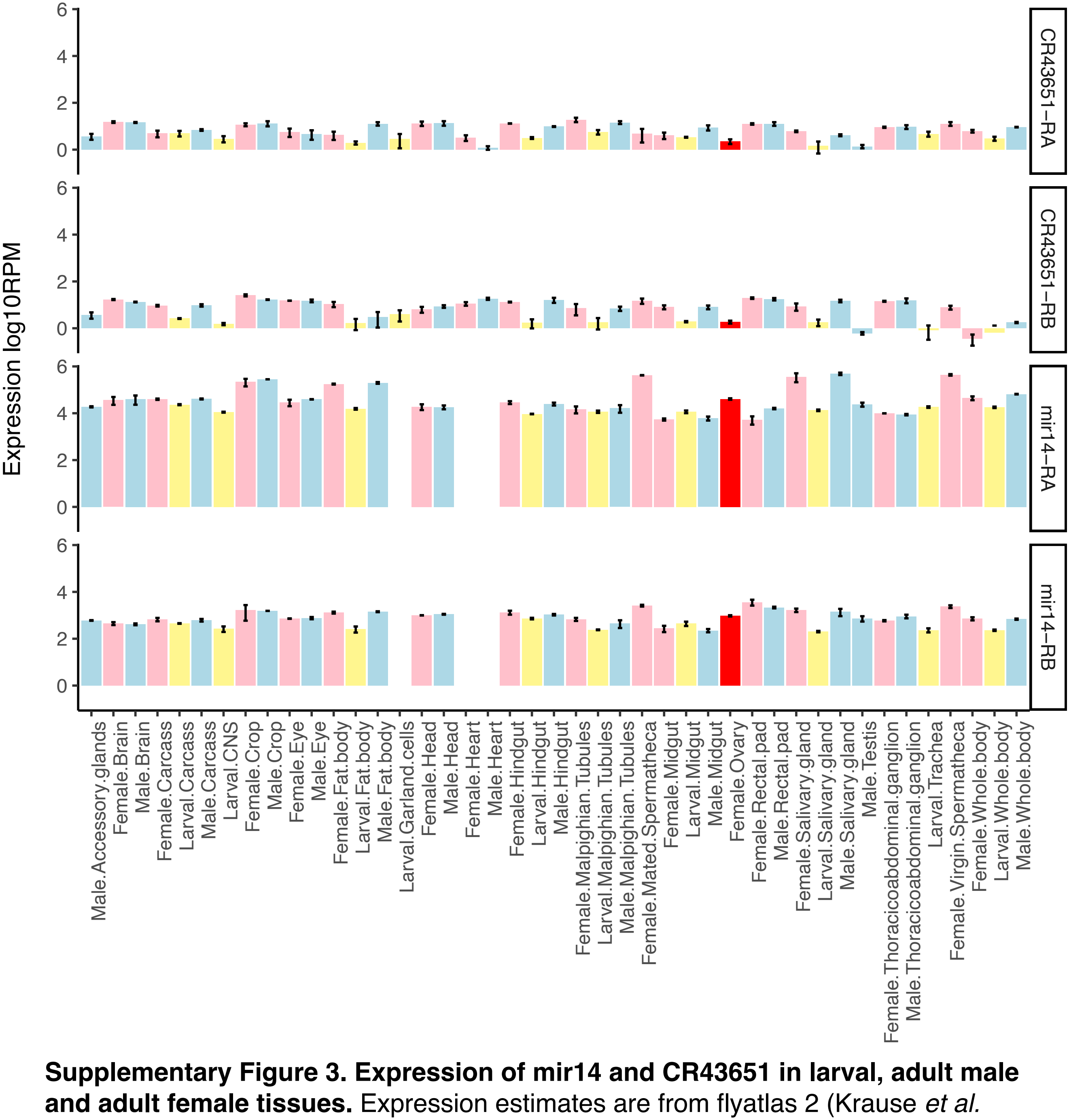


**Supplementary Figure 3. Expression of mir14 and CR43651 in larval, adult male and adult female tissues.** Expression estimates are from flyatlas 2 (Krause *et al.* 2020), and are generated from 2-3 biological replicates of total RNA sequencing (*CR43651*) or small RNA (*mir14*) sequencing. Colors indicate larval tissues (yellow), male tissues (blue) female tissues (pink) and female ovaries (red). Error bars are standard error.
