## Supplementary Figure 1 for "TE invasion fuels molecular adaptation in laboratory populations of *Drosophila melanogaster*"

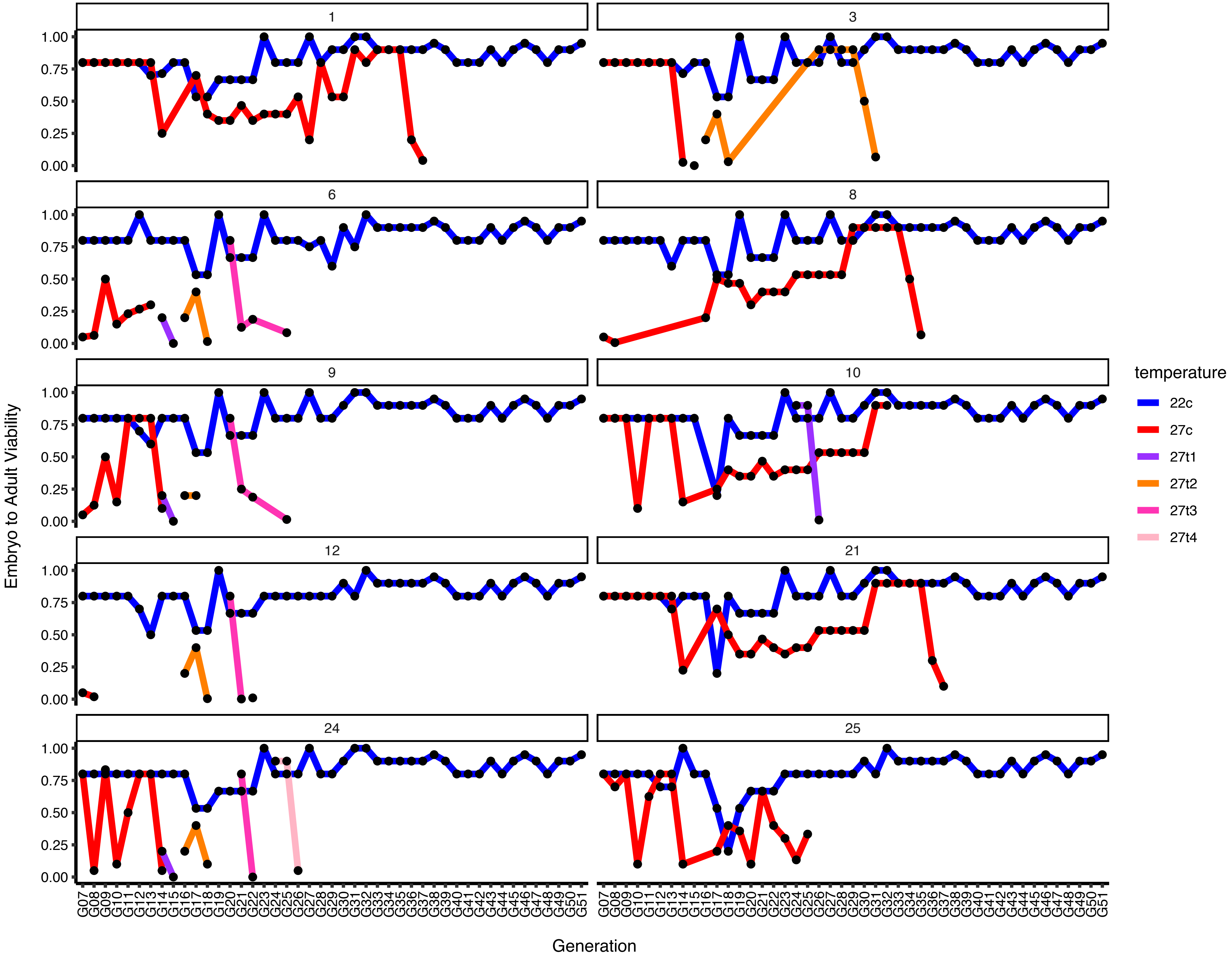


**Supplementary Figure 1. Embryo to adult viability in experimental populations.** Number of embryos was estimated based on volume of embryos collected, and number of adults was rougly counted when each new population was established. Terminated lines at 27C indicate extinction events. Reestablishments of the 27C populations from 22 are represented as different colors.
