## Supplementary Figure 2 for "TE invasion fuels molecular adaptation in laboratory populations of *Drosophila melanogaster*"

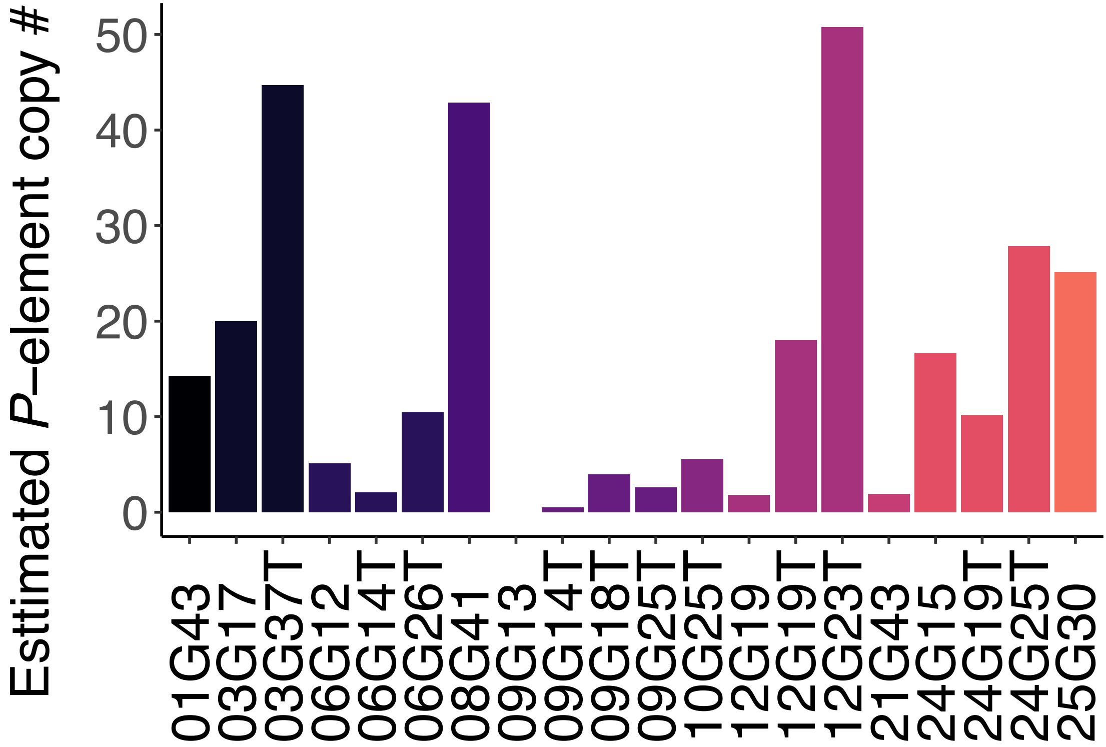


**Supplementary FIgure 2. Estimated P-element copy numbers in 27C populations.** Labels indicate population number first, followed by the generation of extinction. Labels ending in T indicate populations that were restarted from the corresponding sister population reared at 22C. Copy number was estimated by qPCR by normalizing to the single copy gene rpl32, as well as females from the TP5 reference strain (BDSC 64168), which is homozygous for an individual *P*-element insertion.
