## Supplementary Figure 4 for "TE invasion fuels molecular adaptation in laboratory populations of *Drosophila melanogaster*"

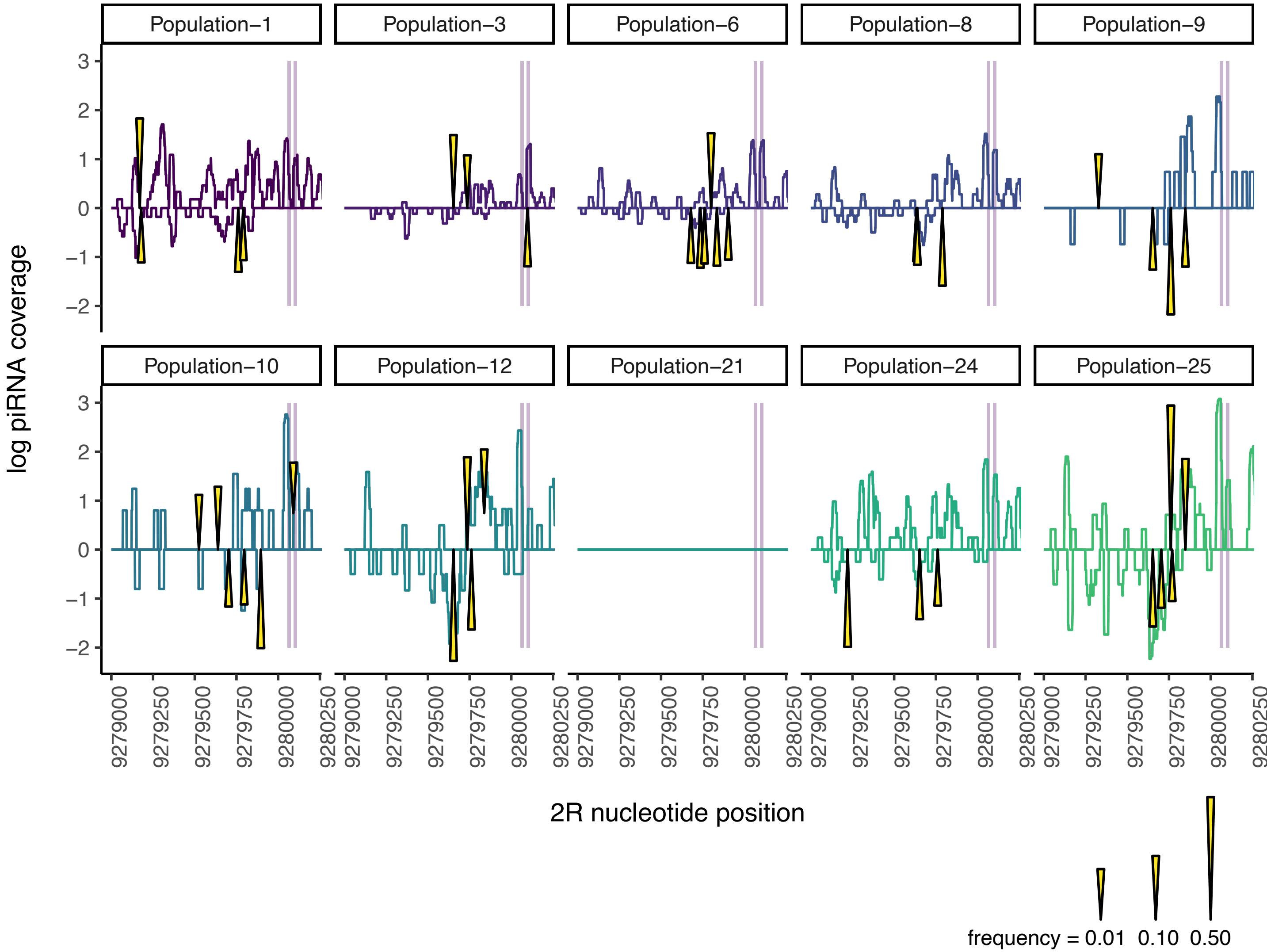


**Supplementary Figure 4. Coincidence of P-element insertions and piRNA production in *lncRNA:43651***. piRNAs from both genomic strands of a portion of *lncRNA:CR43651* represented in Figure 4B. Locations of mir14 transcripts are highlighted in grey. Locations of *P*-element insertions in each population in generation G51 are indicated by yellow triangles, which are scaled according to the estimated insertion frequency. piRNA data are from ovarian libraries generated at Generation 51, abundance is in log-transformed reads per million mapped miRNAs.
