## Supplementary Figure 5 for "TE invasion fuels molecular adaptation in laboratory populations of *Drosophila melanogaster*"

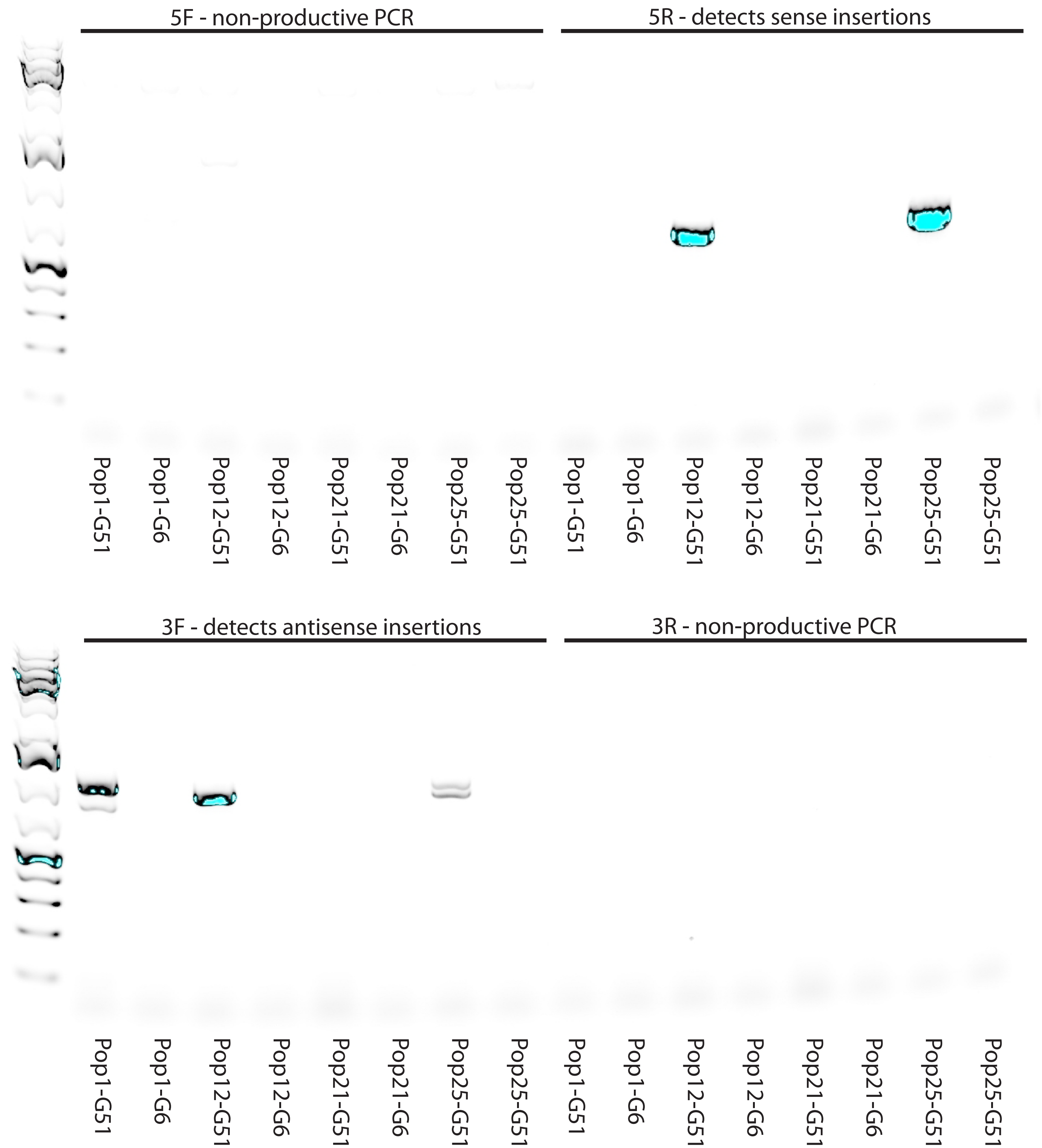


**Supplementary Figure 5. Detection of *lncRNA:CR3651-P*-element chimeric transcripts.** PCRs combining lncRNA:CR43651-F1 with *P*-element-5R and 3F primers detect chimeric transcripts including *lncRNA:CR3651* and sense (5R) or antisense (3F) *P*-element insertions. PCRs using lncRNA:CR43651-F1 and *P*-element 5F/3R fail to produce a bright band due to the large product size (>3 Kb).
