## Supplementary Figure 6 for "TE invasion fuels molecular adaptation in laboratory populations of *Drosophila melanogaster*"

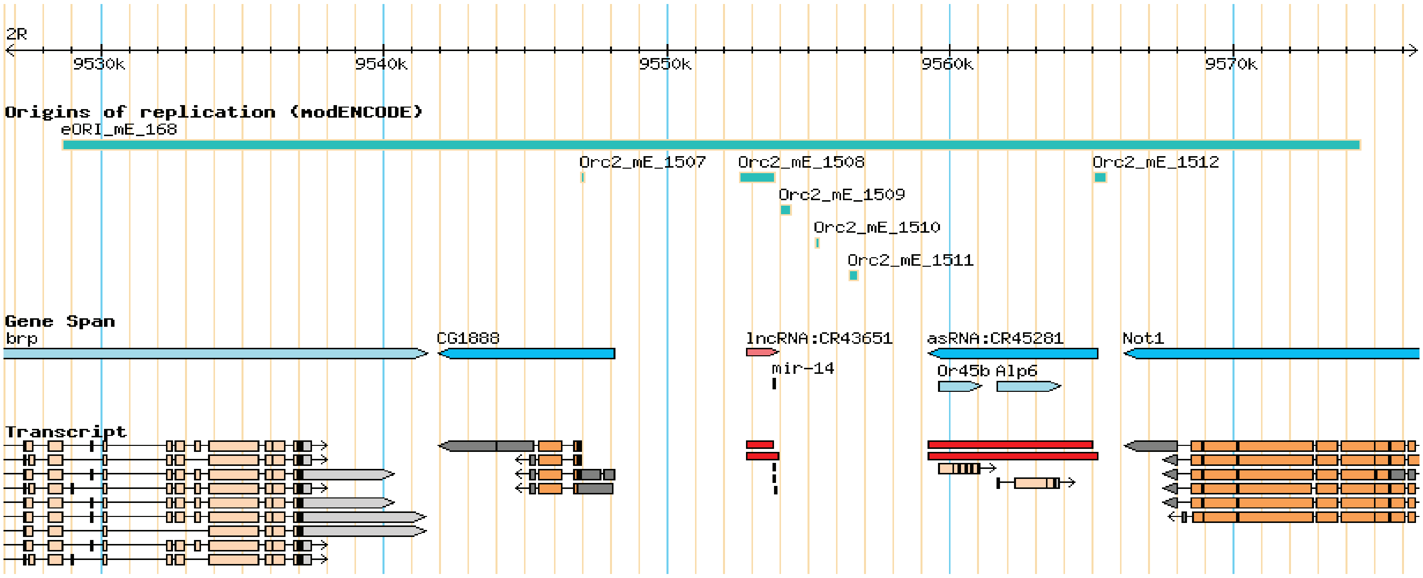


**Supplementary Figure 6. *lncRNA:CR43651* and *mir14* occur in an origin replication.** The genomic window is 2R:9,550,844..9,555,940 from the dm6 reference assembly, rendered in the flybase genome browser.
